# Hepatitis C virus cell culture adaptive mutations enhance cell culture propagation by multiple mechanisms but boost antiviral responses in primary human hepatocytes

**DOI:** 10.1101/2023.11.22.568224

**Authors:** Nicola Frericks, Richard J.P. Brown, Birthe M. Reinecke, Maike Herrmann, Yannick Brüggemann, Daniel Todt, Csaba Miskey, Florian W. R. Vondran, Eike Steinmann, Thomas Pietschmann, Julie Sheldon

## Abstract

Hepatitis C virus (HCV) infection progresses to chronicity in the majority of infected individuals. Its high intra-host genetic variability enables HCV to evade the continuous selection pressure exerted by the host, contributing to persistent infection. Utilizing a cell culture adapted HCV population (p100pop) which exhibits increased replicative capacity in various liver cell lines, this study investigated virus and host determinants which underlie enhanced viral fitness. Characterization of a panel of molecular p100 clones revealed that cell culture adaptive mutations optimize a range of virus-host interactions, resulting in expanded cell tropism, altered dependence on the cellular co-factor micro-RNA 122 and increased rates of virus spread. On the host side, comparative transcriptional profiling of hepatoma cells infected either with p100pop or its progenitor virus revealed that enhanced replicative fitness correlated with activation of endoplasmic reticulum stress signaling and the unfolded protein response. In contrast, infection of primary human hepatocytes with p100pop led to a mild attenuation of virion production which correlated with a greater induction of cell-intrinsic antiviral defense responses. In summary, long-term passage experiments in cells where selective pressure from innate immunity is lacking improves multiple virus-host interactions, enhancing HCV replicative fitness. However, this study further indicates that HCV has evolved to replicate at low levels in primary human hepatocytes to minimize innate immune activation, highlighting that an optimal balance between replicative fitness and innate immune induction is key to establishing persistence.

**Author Summary:** HCV infection remains a global health burden with 58 million people currently chronically infected. However, the detailed molecular mechanisms which underly persistence are incompletely defined. We utilized a long-term cell culture adapted HCV, exhibiting enhanced replicative fitness in different human liver cell lines, in order to identify molecular principles by which HCV optimizes its replication fitness. Our experimental data revealed that cell culture adaptive mutations confer changes in the host response and usage of various host factors. The latter allows functional flexibility at different stages of the viral replication cycle. However, increased replicative fitness resulted in an increased activation of the innate immune system, which likely poses boundary for functional variation in authentic hepatocytes, explaining the observed attenuation of the adapted virus population in primary hepatocytes.

## Introduction

Hepatitis C virus (HCV) establishes a chronic infection in most exposed individuals. Indeed, in 2021, the World Health Organization reported that globally 58 million people were living with a chronic HCV infection [1]. The mechanisms of underlying persistence are complex and remain elusive. One attribute is the high intra-host variability of HCV which is conferred by error-prone replication of the virus. Variants that dominate over time have the survival advantage over those that are lost in a “Darwinian” selection process that results in viral adaptation to specific cellular environments.

There are several lines of evidence that long-term cell culture passage of HCV leads to viral adaptation, conferring optimized usage of host factors in a given cellular context used for viral selection. Interestingly, these adaptive changes in viral phenotypes are often host cell dependent. For example, replication enhancing mutations in subgenomic replicons (SGRs) were shown to compensate for overactivation of the host factor phosphatidylinositol-4-kinase III alpha (PI4KA) in human Huh-7 hepatoma cells [2]. Moreover, ectopic expression of SEC14L2, which has a reduced expression in hepatoma cell lines, is required for the propagation of selected HCV clinical isolates and SGRs *in vitro* but affected replication of cell culture-adapted SGRs only moderately [3, 4].

Previously, Perales and colleagues performed long-term serial passage of the HCV lab strain Jc1FLAG(p7-nsGluc2A) in highly permissive Huh-7.5 human hepatoma cells. After 100 passages, a viral population was recovered with enhanced replicative fitness when compared to the parental virus [5]. The higher fitness of the adapted virus population, named p100 population (p100pop), was associated with increased shutoff of host cell protein synthesis, enhanced phosphorylation of protein kinase R (PKR), and reduced sensitivity to multiple anti-HCV inhibitors [6]. Using p100pop, efficient recapitulation of the entire virus life cycle in stem cell derived hepatocyte like cells (HLCs) was achieved, which was not observed when infecting with Jc1 [7]. Similarly, the enhanced replicative capacity of p100pop also partially overcomes the human-mouse species barrier. Indeed, completion of the full HCV life cycle in humanized murine liver tumor cells and humanized primary mouse hepatocytes upon inhibition of the JAK-STAT pathway, was achieved after p100pop infection but not observed with Jc1 [8]. Genomic sequencing comparison of the p100pop virus population and the parental Jc1 strain revealed fixation of multiple adaptive coding mutations distributed throughout the entire genome [5, 8]. These mutations underlie the improved replication phenotype observed in cell culture, HLCs and engineered murine hepatocytes.

In this study a panel of molecular clones based on the p100pop consensus sequence were generated and phenotypically characterized with the aim of identifying viral and host determinants which enhance HCV fitness. Overall, this study sheds light on the complex mechanisms which allow HCV to establish persistent infection.

## Results

### Genetic characterization of p100pop

Comparison of p100pop consensus sequence with the parental Jc1 genome revealed the emergence of 14 coding mutations distributed throughout the viral genome [5, 8] (Fig. 1A, Table 1). Even though adaptation was performed over 100 passages, mutations were present in the virus population at different frequency levels ranging from around 50 % up to 100 % in the population (Fig. 1B), highlighting the heterogeneity of the adapted virus population. Since the impact of silent mutations for replicative fitness is unclear, we further screened the p100pop consensus sequence for synonymous changes and identified an additional 14 noncoding mutations (Fig. 1A, Table 1). Furthermore, it was noted that the inserted *Gaussia* luciferase reporter gene (Gluc) [9] was partially deleted during the adaptation process, leaving only the foot and mouth disease autoproteolytic peptide sequence (2A) [5], which is subsequently referred to as peptide sequence (PS).

**Fig. 1:**
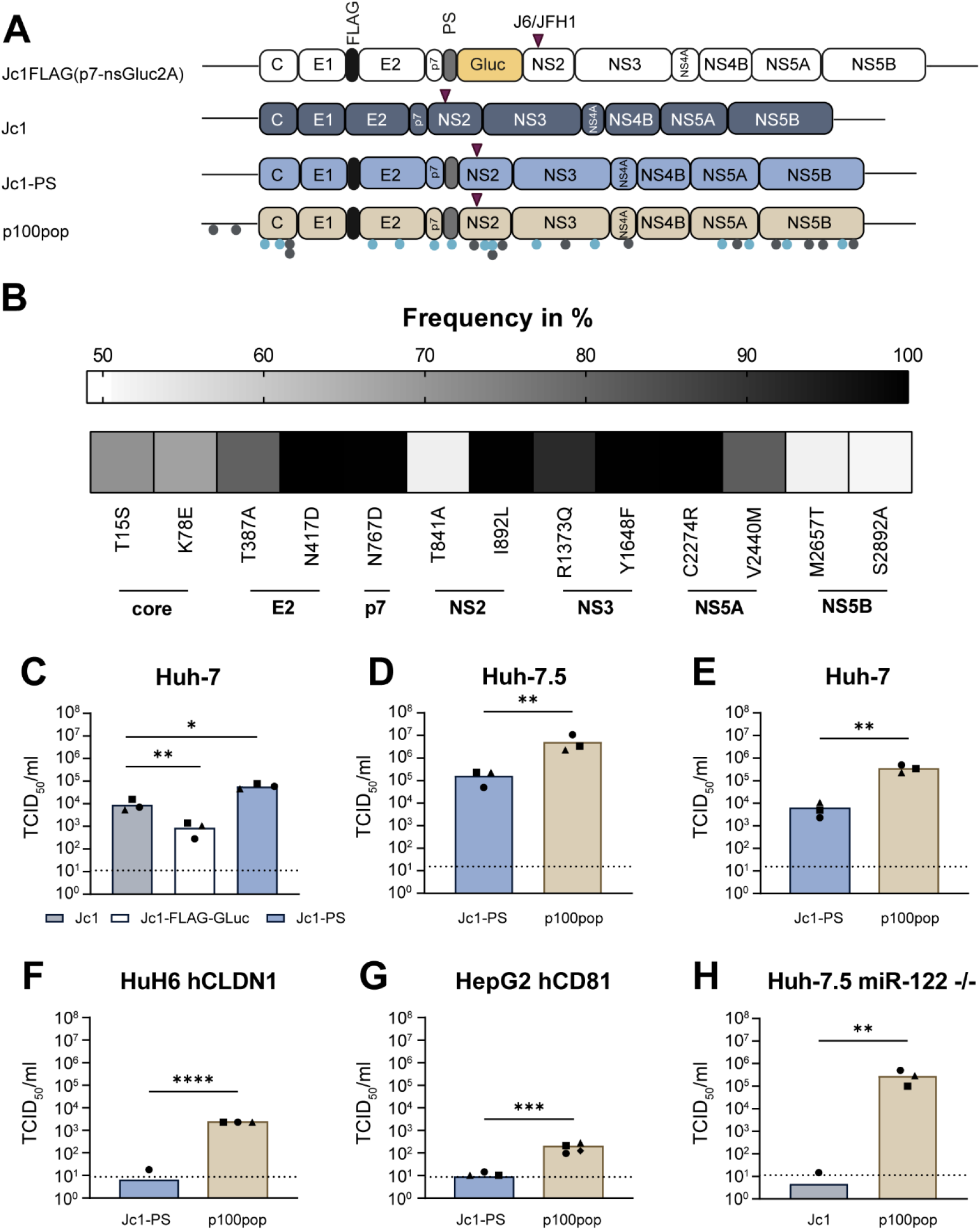
p100pop exhibits enhanced replication fitness in various human liver cell lines. (A) Schematic illustration of the genome organization of the parental virus Jc1FLAG(p7-nsGLuc2A) of p100pop and the Jc1 variants used in this study. Grey dots indicate number and approximate position of noncoding mutations. Light blue dots display coding mutations. The purple triangle reflects the approximate position of the J6/JFH1 junction within the intragenotypic Jc1 chimera. This illustration was generated using BioRender.com. (B) Frequency of adaptive coding mutations detected in the consensus sequence of p100pop. (C) Huh-7 cells were infected with 500 HCV genome equivalents (GE)/cell of different Jc1 variants. (D) Huh-7.5 cells were infected with 10 HCV GE/cell of Jc1-PS and p100pop. (E) Huh-7 cells were infected with 100 HCV GE/cell of Jc1-PS and p100pop. (F) HuH6 cells expressing hCLDN1 were infected with 1,000 HCV GE/cell of Jc1-PS and p100pop. (G) HepG2 cells expressing a HAHA-tagged version of hCD81 were infected with 1,100 HCV GE/cell of Jc1-PS and p100pop. (H) Huh-7.5 miR-122 knockout cells were infected with 140 HCV GE/cell of Jc1 and p100pop. (C-H) After a medium change at 4 h.p.i., cell culture supernatant was collected at 72 h.p.i. and used to determine viral particle production using a limiting dilution assay. Means and individual values of (C-F, H) n = 3 and (G) n = 4 biological independent experiments are shown. Statistical significance was calculated using two-tailed, unpaired t-test with significance levels of p < 0.05 = *, p < 0.01 = **, p < 0.001 = ***, p < 0.0001 = ****. TCID_50_, 50 % tissue culture infectious dose. Dashed lines represent the limit of detection (LOD) of the limiting dilutions assay.

**Table 1:** Mutations in the consensus sequence of p100pop compared to the reference sequence Jc1-PS^a^.

| 5'UTR | Core | E2 | p7 | NS2 | NS3 | NS4A | NS5A | NS5B |
| --- | --- | --- | --- | --- | --- | --- | --- | --- |
| guanine28adenine<sup>b</sup><br>thymine301cytosine<sup>b</sup> | T15S<br>K78E<sup>c</sup><br>G129G<sup>b</sup><br>S173S<sup>b</sup> | T387A<br>N417D | N767D | I830I<sup>b</sup><br>T841A<sup>c</sup><br>G887G<sup>b</sup><br>I892L<br>L900L<sup>b</sup> | R1373Q<br>L1534L<sup>b</sup><br>Y1648F | A1676A<sup>b</sup> | C2274R<br>E2371E<sup>b</sup><br>V2440M<sup>c</sup> | Q2641Q<sup>b</sup><br>M2657T<br>L2834L<sup>b</sup><br>I2874I<sup>b</sup><br>S2892A<sup>c</sup><br>T2979T<sup>b</sup> |

<sup>a</sup> Positions are numbered according to the nucleotide and amino acid coordinates of the infectious clone Jc1-PS (accession number OQ726017) with artificial sequences excluded
<sup>b</sup> Unpublished noncoding mutations identified by NGS
<sup>c</sup> Compared to the Sanger sequencing data novel coding mutations identified by NGS

### Long-term adaptation of HCV broadened the cell culture tropism of HCV

In line with previous reports [10], we could show that the presence of the Gluc results in a significant reduction in viral infectivity which is likely the reason why it is lost during the selection process (Fig. 1C). Based on these observations, we generated a control virus, Jc1-PS, with an identical genome organization to p100pop, containing both the FLAG epitope and PS (Fig. 1A). Interestingly, comparison of viral production rates upon infection of Huh-7 cells between Jc1-PS and unmodified Jc1 revealed a significant increase in viral particle release for Jc1-PS at 72 hours post infection (h.p.i.) (Fig. 1C). Irrespective of this finding, we still detected a significant increase in viral progeny production for p100pop compared to Jc1-PS in Huh-7.5 and Huh-7 cells at 72 h.p.i. (Fig.1D, E). We further investigated the ability of p100pop to replicate in the human liver cell lines HuH6 and HepG2, originating from two different patients diagnosed with hepatoblastoma [11, 12]. Both cell lines are not susceptible to HCV genotype 2a infection unless they ectopically express the human HCV entry factors claudin-1 (hCLDN1) [13, 14] and cluster of differentiation 81 (CD81) (hCD81) [15], respectively. Despite this, permissiveness of these cells for Jc1-PS infection remained low, resulting in infectious particle release below or slightly above the detection limit of the limiting dilution assay (Fig. 1F, G). In contrast, p100pop produced significantly more infectious HCV particles with a titer of around 2.3 × 10^3^ TCID_50_/ml from HuH6 hCLDN1 and 1.7 × 10^2^ TCID_50_/ml from HepG2 hCD81 cells. Whereas the detailed molecular mechanism for limited permissiveness of HuH6 cells is not completely understood, in HepG2 cells this can be attributed to low endogenous expression of the indispensable host factor micro-RNA 122 (miR-122) [16]. To explore whether long-term adaptation modulated dependence of HCV on miR-122, we measured infectious particle release of Jc1 and p100pop in Huh-7.5 cells deficient for miR-122 expression [17] at 72 h.p.i. (Fig. 1H). Confirming our hypothesis, we detected a significant increase in p100pop release from these cells, leading to a virus titer of 3 x 10^5^ TCID_50_/ml in the absence of miR-122 expression, which was not observed with Jc1. In summary, these data demonstrate that long-term adaptation resulted in a host cell-independent increase in replicative fitness of HCV in cell culture, likely partially conferred by a change in miR-122 dependence.

### An infectious consensus clone largely reflects the phenotype of a cell culture adapted HCV population

Phenotypic changes of p100pop correlated with the presence of multiple coding and noncoding mutations in the consensus genome sequence of the virus population [5, 8]. To identify the minimal necessary mutations conferring increased replicative fitness, we generated different infectious p100 consensus clones based on the sequencing data. Here, we focused on the role of coding and noncoding mutations in increased replicative fitness and therefore generated consensus clones harboring either coding (p100NGS_coding) or noncoding mutations (p100NGS_noncoding) alone or a combination of both (p100NGS_all). To investigate the role of adaptive mutations, we excluded the FLAG epitope and the PS from the genome consensus sequence as we previously observed an increase in replicative fitness upon the presence of these artificially introduced sequences (Fig. 1C). This allowed for the use of unmodified Jc1 as a control in the following experiments. Whereas amounts of intracellular viral RNA and progenitor virus remained unchanged for the virus containing noncoding mutations only, we detected a significant increase in replicative fitness upon presence of coding mutations alone or in combination with noncoding mutations (Fig. 2A, B). As differences in replicative fitness between p100NGS_coding and p100NGS_all were minimal, we concluded that adaptive coding mutations are the main driver of enhanced replicative fitness and noncoding mutations contribute only to a minor extent.

**Fig. 2:**
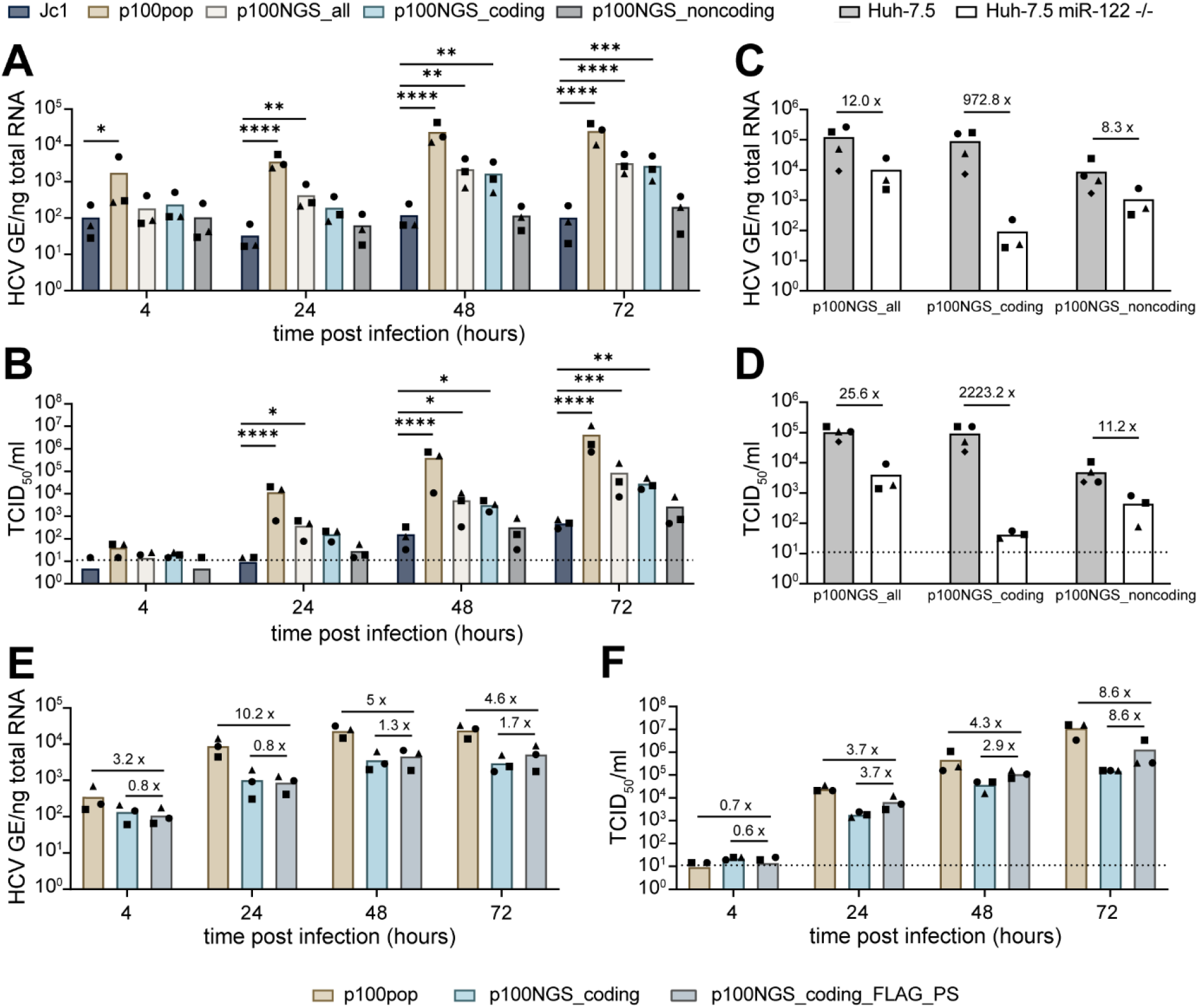
Adaptive coding mutations are indispensable for enhanced replication fitness. (A, B) Huh-7 cells were infected at a MOI of 500 HCV GE/per cell with the indicated molecular clones. Replication fitness was analyzed by (A) RT-qPCR for intracellular viral RNA in total cellular RNA and (B) by virus titer determination of cell culture supernatant at the indicated time points. Statistical significance was calculated using two-way ANOVA with Dunnett’s multiple comparisons test with single pooled variance and significance levels of p < 0.05 = *, p < 0.01 = **, p < 0.001 = ***, p < 0.0001 = ****. (C, D) Huh-7.5 and Huh-7.5 miR-122 deficient cells were infected with 140 HCV GE/cell of the indicated p100 consensus virus variants. At 72 h.p.i. cell culture supernatant and total cell lysates were collected and analyzed for (C) intracellular viral RNA and (D) released infectivity determined by a limiting dilution assay. Numbers indicate the fold-difference of samples collected from Huh-7.5 miR-122 deficient cells compared to samples from the parental cell line. (E, F) Huh-7 cells were infected with 340 HCV GE/cell of p100pop or p100 consensus virus harboring adaptive coding mutation without (p100NGS_coding) or with (p100NGS_coding_FLAG_PS) additional FLAG epitope and PS. Viral fitness was assessed by (E) RT-qPCR for intracellular viral RNA and (F) virus titer determination from cell lysates and cell culture supernatant collected at indicated time points. Numbers indicate fold-difference in viral fitness. Means and individual values in this figure are displayed for n = 3-4 experiments. Dashed lines represent the LOD of the performed limiting dilution assays.

As we observed that long-term adaptation modulated viral dependence on the host factor miR-122, we further investigated replicative fitness of the different consensus clones in Huh-7.5 miR-122 deficient cells. We detected a more than 900-fold difference in intracellular viral RNA and a more than 2,000-fold difference in viral particle release for p100NGS_coding between Huh-7.5 and Huh-7.5 miR-122 deficient cells (Fig. 2C, D). On the contrary, the p100 clone harboring only noncoding mutations replicates at much higher levels in absence of miR-122 expression, reducing the difference only to 8- and 11-fold in intracellular viral RNA and extracellular progenitor virus detected at 72 h.p.i, respectively. These results highlight, that even though noncoding mutations contribute only to a minor extent in cells with abundant miR-122, they are responsible for the observed change in miR-122 usage. Interestingly, combining adaptive coding and noncoding mutations further increases replicative fitness of the p100 clone in the absence of miR-122 expression, leading to amounts of intracellular viral RNA and released infectivity around 10-fold higher than determined for the clone harboring only noncoding mutations (Fig. 2C, D). Together, these observations confirm that adaptive coding mutations are essential for the enhanced replicative fitness of p100pop.

Although presence of adaptive coding mutations significantly increases replication and infectious particle production of the consensus clone, combining with noncoding mutations does not result in a fitness level comparable to the original virus population (Fig. 2A, B). Apart from the missing polyclonal property in the p100 consensus clones, these clones differ in their viral genome structure by the absence of the FLAG epitope and the PS. As we observed a fitness enhancing effect induced by these artificial sequences (Fig. 1C), we next explored to what extent the replication fitness of the p100NGS_coding clone can be improved by the presence of the FLAG epitope between E1 and E2, as well as the PS at the N-terminus of NS2. Therefore, we cloned these sequences into the viral genome of the p100 clone (p100NGS_coding_FLAG_PS) and measured levels of intracellular viral RNA and extracellular progenitor virus in comparison to p100pop and the parental p100NGS_coding clone upon infection of Huh-7 cells. In line with our previous observations, we detected an increase of up to 1.7-fold in intracellular viral RNA and up to 8.6-fold in infectious particles released to the cell culture supernatant for p100NGS_coding_FLAG_PS compared to the parental virus p100NGS_coding (Fig. 2E, F). However, viral replication and particle production of p100pop are still up to 10.2- and 8.6-fold higher than the newly generated clone. In summary, these results show that a molecular clone based on the consensus sequence of p100pop largely reflects the phenotype of the long-term cell culture adapted HCV population.

### Adaptive coding mutations increase virus spread

Based on the observation that coding mutations are largely responsible for the increase in replicative fitness, we hypothesized that selectively advantageous mutations result in an optimization of processes during a defined viral life cycle step. To link the contribution of the p100pop adaptive coding mutations to a distinct viral life cycle step, we generated chimeric constructs with portions of Jc1 and p100 viruses, focusing primarily on proteins involved in entry as well as virus particle assembly and egress (Core-NS2) or proteins involved in viral genome amplification (NS3-NS5B) (Fig. 3A). Neither intracellular viral RNA accumulation nor the amount of released infectious particles were affected by insertion of the p100 coding mutations in the replicase protein sequence of Jc1 (Fig. 3B, C). These data suggest that coding mutations in NS3 to NS5B do not affect processes of genome amplification and consequently release of infectious progeny. To confirm this hypothesis, we generated a p100 SGR including the six adaptive coding mutations located in the nonstructural proteins. RNA replication efficiency of the p100 SGR was assessed in correlation to the firefly luciferase (Fluc) reporter activity upon transfection into Huh-7 cells and compared to the parental JFH1 SGR and a replication incompetent replicon with a deletion in the active site of the viral polymerase (ΔGDD). In line with the presented chimeric full-length virus data (Fig. 3B), transfection of the p100 SGR resulted in relative light unit (RLU) counts similar to the parental JFH1 SGR (Fig. 3D).

**Fig. 3:**
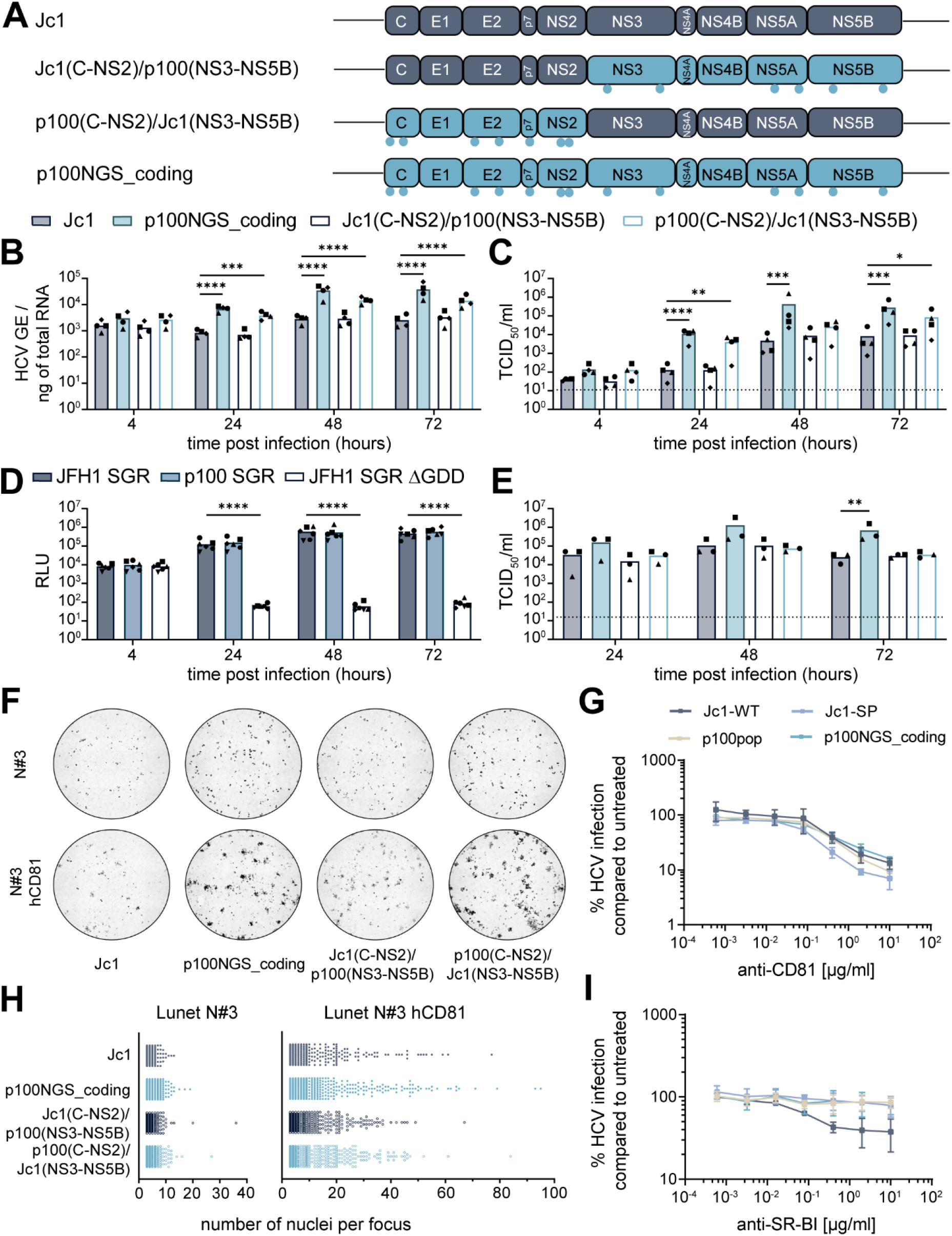
Adaptive coding mutations synergistically improve virus fitness. (A) Schematic illustration of the genome organization of Jc1, p100NGS_coding and the generated Jc1/p100 chimeras. Light blue dots indicate the approximate position of coding mutations present in the viral genomes. This illustration was generated using BioRender.com. (B, C) Huh-7 cells were infected with Jc1, p100NGS_coding and two different Jc1/p100 chimeras with a MOI of 1,500 HCV GE/cell. Replication fitness was analyzed by (B) RT-qPCR for intracellular viral RNA in total cellular RNA and (C) by virus titer determination of cell culture supernatant, both collected at indicated time points. Means and individual values are displayed of n = 4 experiments. (D) *In vitro* transcribed RNA of the p100, JFH1 and a replication inefficient SGR were electroporated into Huh-7 cells. Replication was measured at indicated time points in correlation to Fluc activity of the containing reporter gene. Mean and individual relative light unit (RLU) counts from n=6 experiments are displayed. (E, F, H) *In vitro* transcribed full-length RNA of Jc1, p100NGS_coding and two different Jc1/p100 chimeric viruses were electroporated into Lunet N#3 cells. (E) Virus titers of cell culture supernatant collected at indicated time points were determined using a limiting dilution assay. Means and individual values are displayed of n = 3 experiments. (F, H) Lunet N#3 cells transfected with HCV RNA were mixed with naïve Lunet N#3 with and without hCD81 expression. Cells were fixed at 48 h post electroporation (h.p.e) and stained for NS5A protein expression (black). Microscopic images were taken at 4x magnification and processed using FIJI and CellProfiler. (F) Representative images of whole 96-wells. (H) The number of nuclei per focus were counted automatically using CellProfiler. Only foci with ≥3 nuclei were included in the visualization. Both graphs display all analyzed foci of n=3 experiments, with each individual dot representing one single focus. (G, I) Huh-7.5 cells were treated with serial dilutions of anti-CD81 or anti-SR-BI antibody for 1 hour at 37 °C, before cells were inoculated with the respective virus for additional 3 hours. At 48 h.p.i. cells were stained for HCV NS5A. HCV infection was normalized by calculating the ratio of the mean focus forming units (FFU) per wells treated with the antibody to the mean number of FFU/well of untreated wells. Cells treated with the same concentrations of an isotype control antibody served as control and did not show inhibitory effects on HCV infection. The experiment was done with three technical replicates of treated and untreated cells per biological replicate. Means and ± standard deviations are shown for n = 3 experiments. (B, C, D, E) Statistical significance was calculated using ANOVA with Dunnett’s multiple comparisons test with single pooled variance, p < 0.05 = *, p < 0.01 = **, p < 0.001 = ***, p < 0.0001 = ****. (C, E) Dashed lines represent the LOD of the performed limiting dilution assays.

In contrast, replacement of the Core to NS2 sequences in the full-length chimeric virus resulted in a significant increase in intracellular viral RNA compared to the Jc1 control (Fig. 3B). Furthermore, a trend towards increased amounts of progenitor virus released to the cell culture supernatant was observed (Fig. 3C). To dissect the underlying molecular mechanisms in more detail, we transfected *in vitro* transcribed chimeric full-length virus RNA into Lunet N#3 cells, which are refractory to HCV infection due to low CD81 expression levels [18]. In contrast to the significant differences in infectious titers detected for the p100/Jc1 chimera harboring adaptive mutations in Core to NS2 proteins upon infection of Huh-7 cells (Fig. 3C), the number of infectious particles released to the cell culture supernatant remained unaltered upon transfection of viral RNA into Lunet N#3 cells when compared to Jc1 (Fig. 3E). Losing the fitness enhancing effects of adaptive mutations by transfection of viral RNA into cells with minimal CD81 expression suggests that Core-NS2 mutations enhance virus cell-to-cell spread, as a detectable CD81 expression is required for efficient cell-cell transmission [19]. In order to explore the impact of adaptive mutations on cell-to-cell spread, we mixed HCV transfected Lunet N#3 cells with naïve Lunet N#3 with or without hCD81 expression and analyzed cell-to-cell spread by immunofluorescence-based quantification of the size of the formed foci. Here, we observed not only an increase in the intensity of antigen staining but also an increased number of foci containing greater numbers of nuclei per focus for p100NGS_coding and for the chimeric virus with Core-NS2 mutations (Fig. 3F, H). As interaction of the viral glycoproteins with CD81 and scavenger receptor B-I (SR-BI) is of high importance not only during cell-to-cell transmission but also cell-free entry [19], we further investigated sensitivity of the two Jc1 variants as well as the p100 population and clone to CD81 and SR-BI receptor blockage by antibody treatment. Whereas p100pop and the molecular clone p100NGS_coding were sensitive to CD81 blockage in a manner comparable to Jc1, the modified Jc1 variant, which harbors a FLAG epitope linked to the N-terminus of E2, showed increased sensitivity (Fig. 3G). In contrast, the p100 population and clone showed, similar to Jc1-PS and in contrast to Jc1, lower dependency on the entry receptor SR-BI (Fig. 3I). Taken together, these data imply that adaptive mutations in the structural proteins facilitate virus spread likely due to altered dependence on the entry receptor SR-BI.

The p100NGS_coding clone produced increased amounts of infectious progeny virus compared to Jc1 upon transfection of *in vitro* transcribed RNA (Fig. 3E). As this was observable despite the lack of CD81-dependent cell-to-cell spread, our data further suggest that a combination of multiple mutations, located in structural and nonstructural proteins, cooperatively enhance viral particle assembly and release.

### Adaptive coding mutations confer a change in cyclosporin A susceptibility

The presence of drug resistant mutations in complex quasispecies of RNA viruses represents a major challenge for the antiviral treatment of viral diseases. Indeed, the error-prone replication of RNA viruses provides the basis of a large reservoir of phenotypically distinct but closely related viral genomes which include variants resistant to antiviral treatment without prior drug exposure [20]. To investigate whether p100pop adaptive coding mutations confer resistance to antiviral compounds, we utilized JFH1 and p100 SGRs which encode for the proteins targeted (in-)directly by the four antivirals telaprevir, daclatasvir, IFN-α, and cyclosporin A. We performed dose-response assays at 4 hours post electroporation (h.p.e.) of SGR RNA into Huh-7 and evaluated antiviral activity on replication in correlation to the Fluc reporter activity at 48 h.p.e. Both SGRs were sensitive to treatment with telaprevir, daclatasvir or IFN-α to similar degrees (Fig. 4A, B, C). In contrast, presence of the six adaptive coding mutations resulted in decreased susceptibility of the p100 SGR to cyclosporin A treatment (Fig. 4D). Thus, we conclude that long-term adaptation of HCV not only modulated viral dependence on miR-122 but additionally altered dependence of HCV on the essential host factor cyclophilin A, which is targeted by cyclosporin A.

**Fig. 4:**
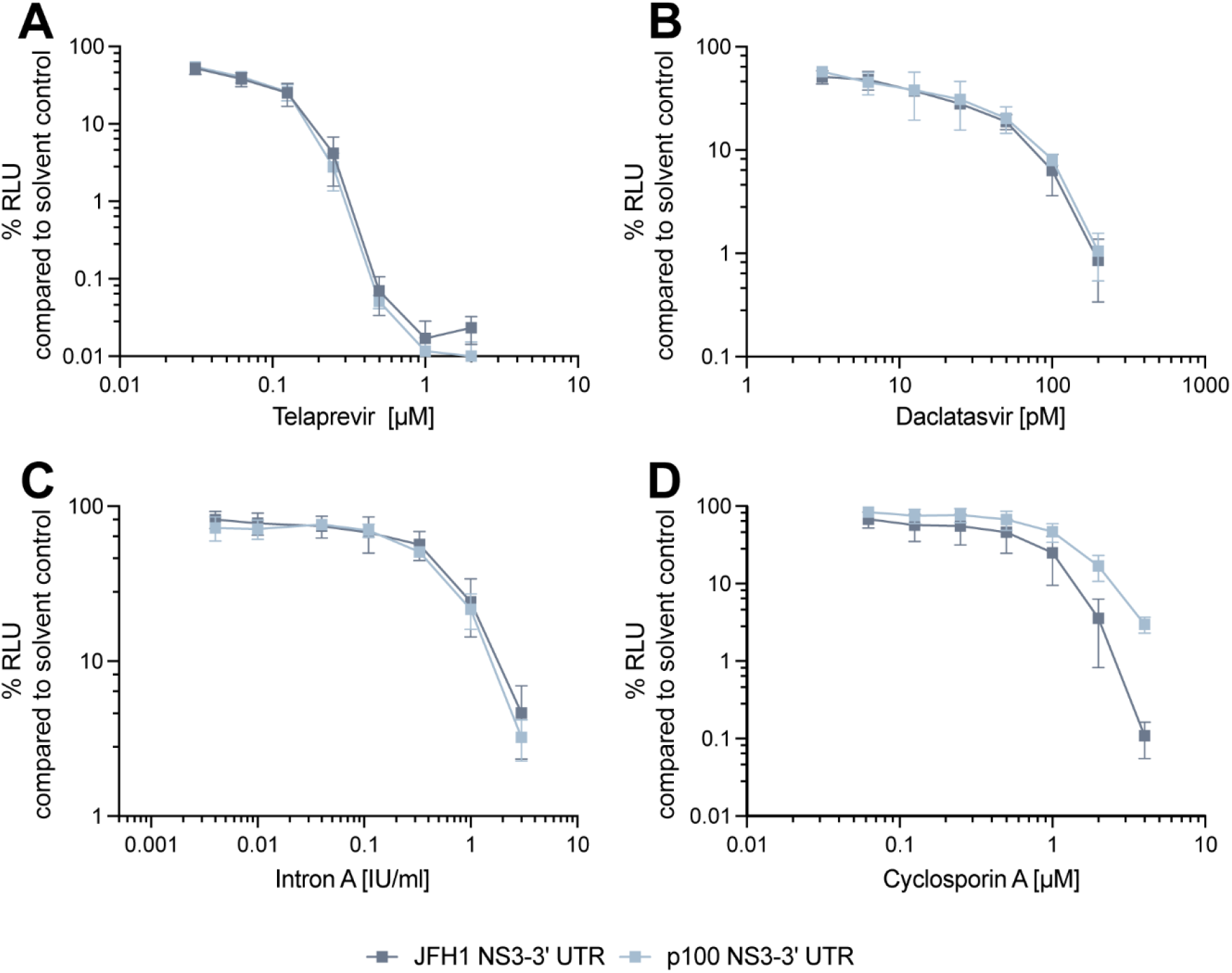
Presence of adaptive coding mutations in the p100 sub genomic replicon results in decreased susceptibility to cyclosporin A treatment. *In vitro* transcribed RNA of the JFH1 and p100 SGR was transfected into Huh-7 cells. At 4 h.p.e. serial dilutions of (A) telaprevir, (B) daclatasvir, (C) Intron A or (D) cyclosporin A were applied. The impact of antiviral treatment on SGR replication was analyzed by measuring Fluc activity at 48 h.p.e. Replication is shown as percentage relative to solvent control-treated cells. Means ± standard deviations are plotted from (A, B, C) n = 3 and (D) n = 4 experiments.

### HCV cell culture adaptation results in viral attenuation in primary human hepatocytes

Since p100pop exhibits enhanced viral propagation rates in several human hepatoma cell lines (Fig.1), we lastly aimed to investigate its replication fitness in primary human hepatocytes (PHHs). A trend towards an increased amount of intracellular viral RNA was detectable for p100pop compared to Jc1-PS at 4 and 24 h.p.i. (Fig. 5A). This correlated with a reduced amount (∼ 1 log) of infectious viral particles released into the cell culture supernatant at 48 and 72 h.p.i. when compared to Jc1-PS (Fig. 5B). In contrast to Huh-7-derived cell lines, PHHs possess intact innate immunity [7, 21].Thus, we hypothesized that elevated amounts of p100pop RNA at early time points induce a greater innate immune response in PHHs when compared to Jc1-PS. As a consequence of this more pronounced activation of the innate immunity, virion secretion at later time points is likely restricted. To confirm this hypothesis, we next explored replication fitness of p100pop in PHH that were (pre-) treated with the JAK/STAT inhibitor ruxolitinib. Whereas the amount of intracellular p100pop RNA was at early time points comparable to the amount measured in untreated PHHs (Fig. 5C), ruxolitinib treatment rescued p100pop particle production rates at 48 and 72 h.p.i. to a level comparable to Jc1-PS (Fig. 5D). These results indicate that cell culture adaptation of HCV results in an attenuated phenotype in PHHs *ex vivo*, likely conferred by a greater induction of the innate immunity.

**Fig. 5:**
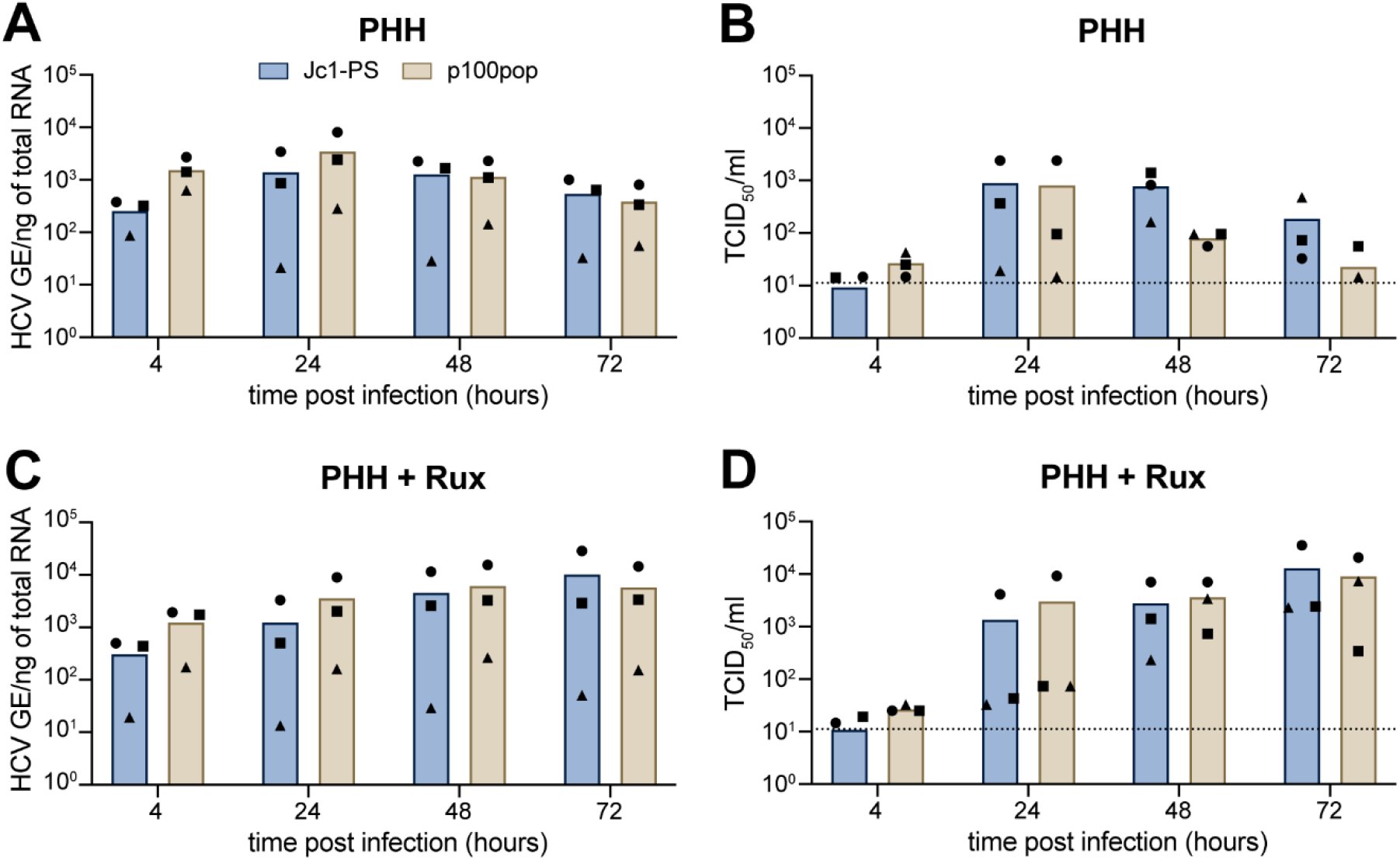
Reduced p100pop particle production of in primary human hepatocytes can be rescued by treatment with the JAK inhibitor ruxolitinib. PHH from three different donors were incubated (A, B) without or (C, D) with 10 µM ruxolitinib (rux) pretreatment. After 16 hours of pretreatment, PHH were inoculated with 100 HCV GE/cell of Jc1-PS or p100pop. At 4 h.p.i. a medium change, including a re-administration of ruxolitinib for pretreated cells, was performed before replication fitness was assessed by (A, C) RT-qPCR for intracellular HCV GE from total cell lysates and (B, D) a limiting dilution assay using cell culture supernatant, both collected at the indicated time points. Means and individual values of infection of three different donors are shown. No statistical significance (p< 0.05) was determined using two-way ANOVA with Sidak’s multiple comparisons test with single pooled variance. (B, D) Dashed lines represent the LOD of the limiting dilution assay.

### Cell culture adaptation results in enhanced innate immune induction in primary human hepatocytes

To explore the molecular mechanisms underlying differences in p100pop fitness between *in vitro* and *ex vivo* cell culture systems, we performed global RNA sequencing (RNA-seq) of infected cells. For this, we inoculated Huh-7 and PHHs with equal amounts of Jc1-PS or p100pop, using conditioned medium (CM) collected from naïve cells as an uninfected control. Total RNA was extracted 28 hours post inoculation (24 h.p.i.) and subjected to RT-qPCR to monitor HCV infection levels. Consistent with previous experiments (Fig. 1E and Fig. 5A), we detected elevated amounts of intracellular p100pop HCV RNA in infected Huh-7 cells and PHHs, when compared to Jc1-PS (Fig. 6A, B). To determine host cell responses to infection with Jc1-PS or p100pop, global transcriptomic analyses were then performed on infected- and conditioned media-treated cells at 24 h.p.i. Statistical analyses of host-cell gene dysregulation identified differentially expressed genes (DEGs) induced upon Jc1-PS and p100pop infection of both Huh-7 (Fig. 6C) and PHH (Fig. 6D). Whereas in Huh-7 cells, the percentage of differentially expressed genes (DEGs) (FDR p-value < 0.05) with a Log_2_ fold change (L2FC) > 1 remains with 0.2 to 0.4 % similar between all tested comparisons (Fig. 6C), the same category of DEGs includes more than 5 % of the analyzed genes in p100pop-infected PHHs and only 2 % in the Jc1-PS-infected PHHs versus CM treatment comparison (Fig. 6D). Interestingly, comparisons of p100pop infection to Jc1-PS in PHHs identified only 6 DEGs with a L2FC > 1.

**Fig. 6:**
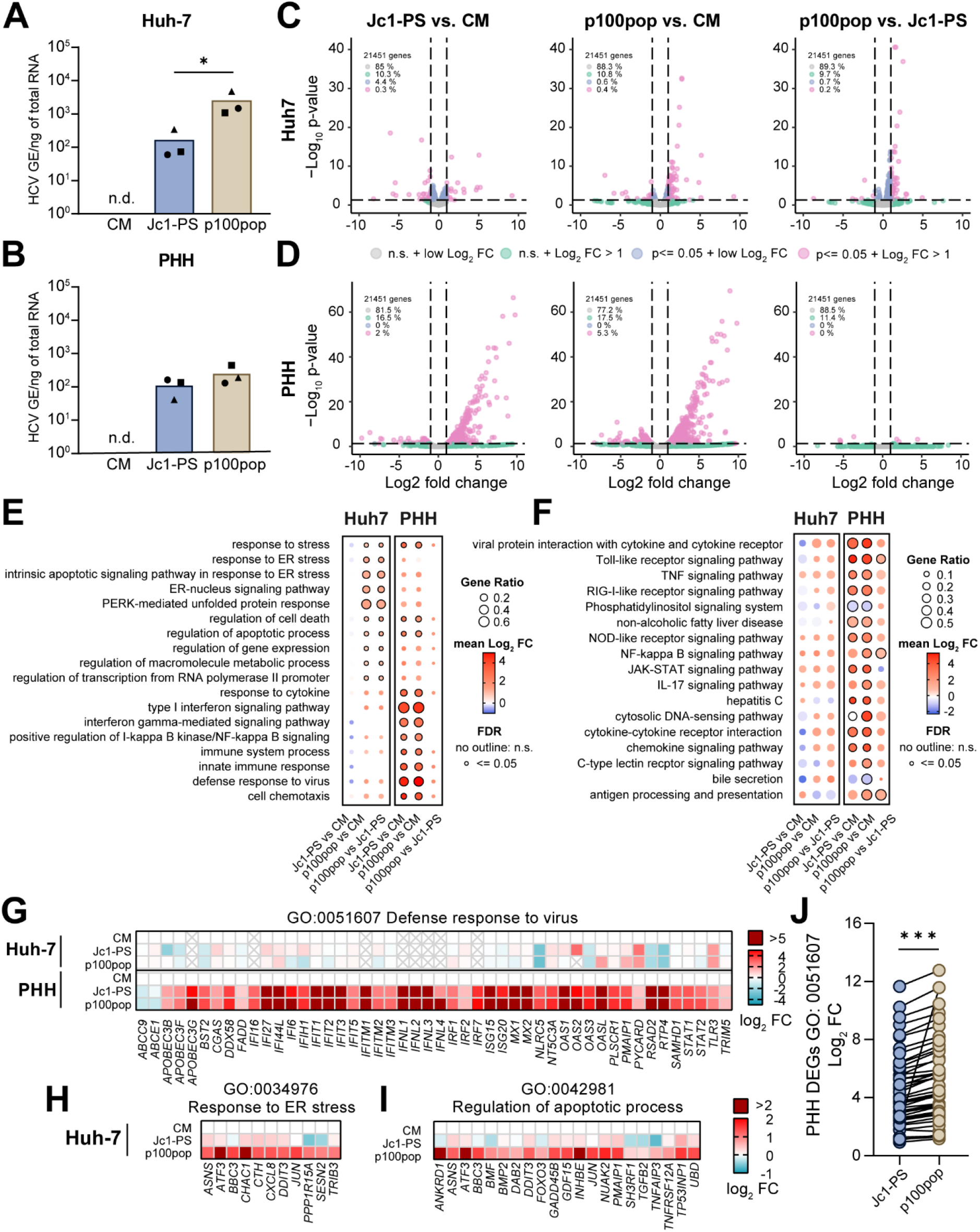
HCV cell culture adaptation results in modulation of host transcriptional responses. Huh-7 and (B) PHH were infected with Jc1-PS or p100pop (MOI 1 TCID50/ml per cell) or incubated with conditioned medium (CM) as an uninfected control. Total RNA was extracted at 24 h.p.i. and used to determine intracellular HCV RNA using RT-qPCR. Statistical significance was calculated using two-tailed, unpaired t-test, p<0.05 = *. (C, D) Volcano plots visualize DEGs induced in infected cells compared to uninfected cells, for Jc1-PS- and p100pop-infected Huh-7 cells (C) and PHH (D). (E) Dot plot visualizes GO enrichment analysis of virus-induced DEGs upon infection of Huh-7 cells (left) and PHH (right). (F) Dot plot visualizes KEGG pathway analysis of virus-induced DEGs upon infection of Huh-7 cells (left) and PHH (right). (G) Heat map visualizing dysregulation of genes (mean log2 FC) associated with the GO category ‘defense response to virus’ upon infection of Huh-7 cells (top panel) or PHHs (bottom panel) with Jc1-PS or p100pop. (H, I) Heat maps visualizing mean log2 FC in gene expression relative to CM-treated cells for genes associated with the GO categories (H) ‘response to ER stress’ and (I) ‘regulation of apoptotic processes’ upon infection of Huh-7 cells with Jc1-PS or p100pop. (J) Comparison of the mean log2 FC of genes clustering in the GO category ‘defense response to virus’ upon infection of PHHs. Statistical significance was calculated using two-tailed, paired t-test, p< 0.001 = ***. (A-I) Data presented derived from infection n = 3 biological replicates. FC, fold change. N.s., not significant.

Infection-induced transcriptional dysregulation modulates a range of cell intrinsic processes and canonical signaling pathways. To investigate these in more detail, we compared host biological processes targeted by the two viruses in both cell systems by performing Gene Ontology (GO) enrichment analyses and KEGG pathway analyses on infection-induced DEGs (Fig. 6E, F). GO categories associated with endoplasmic reticulum (ER) stress, regulation of apoptosis and cell death, as well as the unfolded protein response were significantly enriched upon infection of Huh-7 cells with p100pop, which was not seen in Jc1-PS-infected cells (Fig. 6E). In contrast, in PHH, enriched GO categories were generally associated with innate immune response, viral defense and IFN signaling, regardless of the infecting virus strain. Supporting our GO analyses, KEGG pathway analyses revealed that innate immune signaling pathways are significantly upregulated in PHHs, but not in Huh-7 cells upon virus infection (Fig. 6F). Indeed, visualization of gene expression changes for GO category 0051607 (defense response to virus) highlights pronounced differences in host reactivity to HCV infection between the two cell systems used in this study (Fig. 6G). Innate immunity signatures dominate in PHH but are completely absent in Huh-7 cells. To focus on differences between the two virus strains, we compared the strength of the changes in gene expression by comparing the fold expression changes of genes of the same category. Here, both viruses induce a comparable antiviral defense response upon infection of PHH (Fig. 6G), however infection with p100pop results in a significantly increased induction compared to infection with Jc1-PS (Fig. 6J). These data support our hypothesis that p100pop attenuation in PHHs results from a stronger innate immune responses exerted by the host cells, proportional to the observed increased viral replication.

Additionally, we compared expression of genes from the GO categories 0034976 (response to ER stress) and 0042981 (regulation of apoptotic process) upon infection of Huh-7 cells. Whereas minimal dysregulation was detectable upon Jc1-PS infection, significant up-regulation of the genes involved in regulating processes during ER stress and apoptosis was observed upon p100pop infection (Fig. 6H, I). Thus, our RNA-seq data indicate that infection with p100pop results in a strong and early induction of ER stress in Huh-7 cells, which may contributes to the observed enhanced replication fitness.

## Discussion

The complex molecular mechanisms by which viruses establish persistence are still under investigation but are tightly linked to natural selection of viral genomes with enhanced virus fitness. In this study, we investigated molecular determinants, which confer elevated fitness to a cell culture adapted HCV in order to identify principles by which HCV optimizes its replicative fitness *in vitro*.

Long-term passaging of the HCV strain Jc1FLAG(p7-nsGluc2A) resulted in increased viral fitness of the virus population p100pop in Huh-7, Huh-7.5 and Lunet cells compared to its parental virus [6]. In line with these observations, we could show that p100pop exhibits not only increased infectious viral particle release in Huh-7-derived cell lines (Fig. 1D, E), but also in the hepatoma cell lines HuH6 and HepG2 originating from a different donor and ectopically expressing hCLDN1 or hCD81, respectively (Fig. 1F, G). As Huh-7 and especially the Huh-7.5 subclone are the most permissive cell lines compared to other hepatoma cells, likely due to their host genetic background (high CD81 expression levels, defects in interferon induction) [22, 23], it is interesting that long-term adaptation to Huh-7.5 cells resulted in a host cell-independent increase in virus fitness. HepG2 cells only weakly support HCV replication due to a lack of miR-122 expression [16]. However, infection with p100pop results in a robust release of infectious viral particles from HepG2 hCD81 cells (Fig. 1G). This observation correlates with the presence of the mutation G28A in the 5’ UTR of the p100pop consensus sequence (Table 1), which was previously described as a miR-122 inhibitor resistance associated variant [17, 24]. It was shown that a change from guanine to adenine at this position leads to folding of the HCV internal ribosomal entry site in the 5’ UTR in absence of miR-122/Ago2 binding, which consequently allows virus translation and replication in absence of miR-122 expression [17, 25–27]. In line with this, we detected viral titers up to 3 × 10^6^ TCID_50_/ml upon p100pop infection of Huh-7.5 cells deficient for miR-122 expression (Fig. 1H), implying that the host cell line-independent increase in replicative fitness is likely due to changes in host factor usage, as shown for the miR-122.

Interestingly, infection of PHH resulted in a moderate attenuation of p100pop infectious particle release (Fig. 5B). This observation correlated with an increase in detectable intracellular viral RNA at early time points (Fig. 5A, 6B) and stronger induction of an antiviral defense response compared to the non-adapted virus (Fig. 6J). In contrast, activation of the innate immune response was not detectable in infected Huh-7 cells, regardless of which viral variant was used (Fig 6E, F, G). Consequently, our data highlight that the restriction of HCV infection in PHH, in contrast to Huh-7 cells, is conferred by an early induction of antiviral gene expression. In line with this, transcriptomic analyses of Tegtmeyer and colleagues recently showed that the transcriptional landscape of HCV-infected PHH is dominated by induction of antiviral interferon regulated gene expression, which was not observable in Huh-7.5 cells [21]. Building on this observation, treating PHH prior to and during HCV infection with the JAK/STAT signaling pathway inhibitor ruxolitinib led to higher levels of HCV infection at late time points for both viruses (Fig. 5C, D). However, the numbers of p100pop infectious particles detected in the supernatant remained comparable to Jc1-PS. Other reports provide evidence that elevated basal expression of IRF1 confers early antiviral responses of PHHs by mediating higher basal expression levels of IRF3 activators and interferon regulated genes independent of IFN signaling [21, 28, 29]. In accordance with this, we detected higher expression values of IRF1 in our PHH data set than in the Huh-7 data set (data not shown). Furthermore, a very recent publication revealed a so far uncharacterized antiviral function by PKR, which blocks infectious particle assembly by activating IRF1 [30]. Based on the observation that infection with the long-term cell culture adapted HCV results in enhanced phosphorylation of PKR [6], we propose that attenuation of p100pop viral particle formation is conferred not only by IFN-dependent effector protein expression but also by increased IRF1 activation in PHHs.

Furthermore, we detected a difference in the host cellular response upon infection of Huh-7 cells with p100pop compared to Jc1-PS. Mainly DEGs clustering in pathways associated with ER stress signaling and unfolded protein response were modulated by p100pop (Fig. 6E). The virus-induced stress response results in halt of host protein translation due to elevated levels of phosphorylated eIF2α [31], which was previously described to be enhanced upon p100pop infection [6]. Inhibition of antiviral effector protein synthesis as well as stress granules formation was described to be pro-viral for HCV infection [32–34]. Additionally, ER stress induces autophagy which leads to formation of double-membrane vesicles called autophagosomes. Previous publications provide evidence that HCV replication can take place on membranes of these vesicles and that essential autophagy related proteins are hijacked by HCV for its own replication and translation [35–39]. In line with this we detected an enhanced upregulation of *DDIT3* and *SESN2* upon p100pop infection of Huh-7 cells (Fig. 6H), which were previously described to induce autophagy [40–42]. These results suggest that early induction of the ER stress response is not only a consequence of enhanced fitness but may possibly also contributes it.

In order to dissect the molecular mechanisms underlying the enhanced fitness in more detail, we generated different molecular p100 clones, harboring either all adaptive noncoding or coding mutations identified in the p100pop consensus sequence alone or in combination. Replication kinetics experiments revealed that coding mutations are the minimal necessary mutations conferring enhanced replication fitness (Fig. 2A, B). However, replication fitness of the molecular clones p100NGS_coding and p100NGS_all does not reach fitness levels comparable to the virus population. One additional difference from the polyclonal property is the removal of artificial sequences present in the viral genomes of the quasispecies. Interestingly, we observed that, in contrast to the insertion of the full Gluc reporter, insertion of the FLAG epitope and the PS into the Jc1 backbone resulted in an increase of detected infectivity in the cell culture fluid (Fig. 1C). Additionally, insertion of the FLAG epitope and PS into the p100NGS_coding clone resulted in an increased amount of intracellular viral RNA as well as infectious progenitor virus detected in the cell culture supernatant (Fig. 2E, F). Therefore, we hypothesize that linking of the FLAG epitope to the N-terminus of E2 leads to conformational changes, which alter receptor dependence as documented by enhanced sensitivity of the FLAG-tagged Jc1 to an CD81-targeting antibody and more resistance of the same virus to an antibody against SR-BI (Fig. 3G, I).

However, it is important to note that the impact of the artificial sequences is more pronounced on the amount of infectious virus particles detected in the cell culture fluid than intracellular viral RNA (Fig, 2E, F). Apart from the FLAG epitope, also the PS could possibly contribute to enhanced virus fitness. In contrast to the unmodified Jc1, where the junction site between p7 and NS2 is cleaved by the signal peptidase, NS2 is released by auto cleavage upon presence of the PS. Consequently, differences in cleavage kinetics can affect downstream processes that further influence virus fitness as it was shown for the mutation V2440L and V2440A in NS5A, affecting NS5A-NS5B cleavage kinetics and fitness of HCV variants [43, 44]. Notably, NGS revealed the presence of a single coding mutation located in the PS consensus sequence of p100pop (Fig. 1A) which was not investigated but could contribute to enhanced fitness of p100pop as well. However, even in presence of adaptive coding mutations and the FLAG epitope and the PS, the molecular p100 clone still exhibits viral replication and particle production up to 10-fold lower than the virus population (Fig. 2E, F). Since the frequencies of the adaptive coding mutations varies within the viral population (Fig. 1B), it is possible that the consensus sequence differs from the master sequence, which is the dominant sequence built by the most fit viral variants within the quasispecies [45], and therefore only partially reflects the phenotypic changes gained during long-term cell culture adaptation.

To directly examine the mechanism(s) of adaptation, we used the molecular clone p100NGS_coding and explored on which life cycle step adaptive coding mutations identified in the consensus sequence confer fitness enhancing effects. For this, we generated chimeric viruses harboring either structural proteins, p7 and NS2 (C-NS2) or only nonstructural proteins (NS3-NS5B) of the p100NGS_clone (Fig. 3A). With this approach, we aimed to distinguish between fitness enhancing effects targeting viral replication, or processes during virus entry and virus particle assembly and egress. We observed fitness levels comparable to Jc1 for the chimeric virus harboring the replicase complex from the p100 clone (Fig. 3B, C). This was unexpected as this chimera includes a previously described titer enhancing mutation in NS5A, C2274R [46, 47]. However, we also identified V2440M in NS5A as a second adaptive mutation in NS5A. As mentioned earlier, position V2440 is tightly linked to the fitness of viral variants. Whereas a valine to leucin exchange resulted in a fitness increase, an alanine at the same position strongly impaired viral fitness [43, 44]. The impact of V2440M on virus fitness remains to be explored but it is possible that C2274R became fixed in order to compensate for deleterious effects of V2440M to reach wild-type fitness levels. Unchanged levels of intracellular viral RNA, together with p100 SGR replication comparable to the JFH1 SGR (Fig. 3D), indicate further that adaptive mutations in the nonstructural proteins do not influence efficiency of viral genome amplification or likely also not the formation of replication organelles. This is in consent with previous published literature: Replication enhancing mutations often alter phosphorylation levels of NS5A [48, 49], which confers an imbalance between viral genome amplification and production of progenitor virus [50–52] and are therefore unlikely to be selected. We observed a significant increase in intracellular viral RNA and progenitor virus release upon mutating mainly structural proteins (Fig. 3B, C). Transfection of *in vitro* transcribed RNA of this clone into Lunet N#3 cells resulted in virus particle release comparable to Jc1 (Fig. 3E). Lunet N#3 cells express only low levels of CD81 and are therefore refractory to cell-free HCV infection [18] and likely allow also only low levels of cell-to-cell spread as seen for CD81-negative HepG2 cells [53]. Based on this, our observations in Fig. 3E hint towards a role of adaptive Core-NS2 mutations in enhanced virus spread. This hypothesis is further promoted by the detection of an increased size in viral foci for the chimeric virus p100(C-NS2)/Jc1(NS3-NS5B) (Fig. 3F). Virus spread via direct cell-to-cell transmission remains unaffected by presence of neutralizing antibodies and therefore plays an important role in the establishment of persistent infection [53]. Glycosylation of the viral envelope is another key mechanism of HCV to evade from neutralizing antibodies during cell-free transmission [54]. Here, N417 is an important glycosylation site in the glycoprotein E2, shielding the virus from neutralizing antibodies [55]. Mutational changes at amino acid position 417 are often found in viruses adapted to cell culture due to absence of selection pressure exerted by neutralizing antibodies [56, 57]. Increased exposure of neutralizing epitopes was previously reported to correlate with an increase in viral fitness likely due to increased accessibility of the CD81 binding site [57]. However, neither p100pop nor the p100 molecular clone, both harboring an adaptive coding mutation at position 417 (Table 1), showed an altered sensitivity to blocking of the HCV receptor CD81 compared to Jc1 (Fig. 3G). In contrast, both viruses exhibited low sensitivity to a pretreatment with an anti-SR-BI antibody (Fig. 3I), leading to the hypothesis that increased virus spread of these viruses is conferred by decreased SR-BI dependency.

Our data further indicate that multiple mutations cooperatively contribute to the enhanced fitness of the molecular clone p100NGS_coding since the combination of all mutations in this molecular clone resulted in the most pronounced change in replicative fitness (Fig. 3B, C, E). The increased accumulation of infectivity (Fig. 3C, D) can be explained by multiple hypothesis: Even though a more vigorous accumulation of intracellular RNA was detected, our SGR data (Fig. 3D) suggest that it is unlikely that the more rapid accumulation of viral RNA permits production and accumulation of a greater quantity of infectivity. It is thus tempting to speculate that processes of viral particle assembly or release are likely to be more efficient by presence of adaptive mutations. HCV viral particle assembly is complex and mechanistic details remain elusive. However, they very likely assemble in close proximity to lipid droplets [58]. A cell culture adaptive mutation in p7 of JFH1, C766Y, which is adjacent to our identified mutation N767D, was shown to increase the size of lipid droplets and thereby likely contributes to enhanced fitness of the analyzed cell culture adapted virus [59]. A similar mutational change from asparagine to aspartic acid was found at position 765 in p7 of an adapted JFH1-based virus, which also correlated with enhanced virus progenitor release [44]. Even though, we did not further analyze the role of N767D in enhanced virus fitness, it is possible that mutations at this position modulate lipid droplet morphogenesis and thereby provide an improved platform for interactions of viral proteins involved in viral particle formation. Interaction of p7 and NS2 is also essential for virus particle assembly, as both together coordinate the shuttling of E1E2 heterodimers as well as HCV Core to the site of virus assembly [60–62]. In cell culture adapted intra- and intergenotypic HCV chimeras, mutations in p7 and NS2 are frequently found and likely compensate for incompatibilities between those proteins, resulting in enhanced viral particle production [60, 63]. Further, it is known that NS5A and Core colocalize at lipid droplets as an intermediate step [64, 65], with NS5A likely transporting newly generated genomic RNA to the site of assembly for nucleocapsid formation. The adaptive coding mutation T15S is located in the RNA-binding domain of Core [66] and could therefore be important for the transfer of nascent genomic RNA from NS5A to Core. Overall, our observations together with previously published work highlight that optimization of the complex interactions of viral proteins during the assembly process is one key determinant of enhanced fitness of cell culture-adapted HCV.

Lastly, we analyzed the role of adaptive coding mutations in the broad antiviral resistance of p100pop [6]. For this, we analyzed sensitivity of the p100 SGR to different classes of anti-HCV treatment. Interestingly, we observed that the six adaptive coding mutations confer a partial resistance against inhibition of cyclophilins, but not against the other tested antivirals (Fig. 4). Results reported by Sheldon and colleagues showed that biological clones of the population, obtained by limiting dilution, exhibit a replication fitness and telaprevir resistance similar to the parental population [6]. In contrast, the p100 SGR exhibits neither an increased replication fitness (Fig. 3D) nor resistance against telaprevir, daclatasvir and IFN treatment (Fig. 4A, B, C). Thus, our and their results suggest that HCV cross-resistance against different classes of antiviral treatment is conferred by a general increase in viral fitness, being sufficient for viral adaptation to selection pressure exerted by antiviral treatment, and not by an accumulation of resistance associated mutations during the extended time of infection.

In contrast, the presence of adaptive coding mutations in NS3-NS5B resulted in a partial resistance against cyclosporin A treatment (Fig. 4D). The main target of cyclosporin A is cyclophilin A (CypA), which is an important co-factor for HCV infection [67]. Interaction of NS5A with CypA via its isomerase active site promotes RNA-binding properties of NS5A and thereby the formation of the replication complex with NS5B [68–71]. Interaction of CypA with NS5B in a ternary complex with NS5A further regulates genome amplification [72]. Additionally, formation of replication organelles was shown to be abrogated by cyclophilin inhibition [73–75]. Mutations at position 2440 in NS5A (V2440L/A) were not only associated with virus fitness in general (see above) but also with partial cyclosporin A resistance [43, 44]. Interestingly, V2440L was detected at a low frequency within the quasispecies of the parental virus population and the population at 45 passages [6], before the exchange to methionine was fixed in the consensus sequence of p100pop with around 80 % (Fig. 1B). If V2440M modulates virus fitness and cyclosporin A sensitivity by delayed processing of the NS5A-B junction, as previously proposed for V2440L and V2440A [43, 44], remains unclear. In addition, mutation C2274R is of high interest in regards to cyclosporin A resistance as it is located within the PKR binding domain of NS5A [76]. Recent work suggests a regulatory function of CypA in complex with NS5A on PKR-dependent antiviral activity during HCV infection [75]. Sheldon and colleagues further showed that enhanced replication fitness of p100pop correlated with increased phosphorylation of PKR [6], promoting our hypothesis that long-term adaptation resulted in an altered interaction between HCV and CypA and thereby possibly also influencing PKR activity. However, in depth investigation of the mechanistic details is still required.

In summary, this work revealed multiple molecular determinants of enhanced replication fitness of a long-term cell culture adapted HCV. It highlights that increasing fitness of HCV is tightly linked to the optimization of the viral interaction with the host and its environment.

## Material and methods

### Viruses and subgenomic replicons

The generation of the long-term passaged virus population p100pop was previously reported [5, 6]. The infectious clones Jc1 (pFK-i389_JFH1/J6/C-846_dg) and Jc1FLAG(p7-nsGLuc2A) were previously described [9, 77]. The Jc1-PS plasmid (accession number OQ726017) was generated by replacing the complete GLuc sequence with a gBlock fragment (IDT) containing the foot and mouth disease autoproteolytic peptide sequence (2A) from the p100pop consensus sequence using restriction enzymes NotI and BsaBI. The p100NGS_coding plasmid (accession number OQ726016), harboring adaptive coding mutations identified by sequencing [8, 78], was generated by site directed mutagenesis and restriction-based cloning using the Jc1-PS plasmid as a vector backbone. Sequences of the molecular clones p100NGS_all (accession number OQ726014) and p100NGS_noncoding (accession number OQ726013) were designed using the p100pop consensus sequence (accession number KC595609.1, [8]. The sequences were commercially synthesized and cloned into a pUC57 vector (GenScript). The plasmid p100NGS_coding_FLAG_PS (accession number OQ726015) was generated by transferring an insert, containing the E1-NS2 sequence of the p100pop consensus sequence which is flanked by the restriction sites BsiWI and NotI and includes only coding mutations, from a commercially synthesized pUC57 vector into the p100NGS_coding plasmid. The chimeric plasmid p100(C-NS2)/Jc1(NS3-NS5B) (accession number OQ726017) was cloned by digesting of p100NGS_coding plasmid DNA and pFK-i389_JFH1/J6/C-846_dg with the restriction enzymes EcoRV and AvrII. The Jc1(C-NS2)/p100(NS3-NS5B) plasmid (accession number OQ726019) was cloned by transferring inserts from pFK-i389_JFH1/J6/C-846_dg and Jc1-PS into the backbone p100NGS_coding using restriction enzymes BsiWI and AvrII as well as NcoI and BsiWI, respectively.

The JFH1 NS3-3’ UTR SGR (pFK_i389_F-Luc_EI_NS3-3_JFH1) is based on the previously published Con1 SGR plasmid vector [79] with an additional replacement of the neomycin resistance with a Fluc reporter gene. The replication incompetent JFH1 NS3-3’ UTR ΔGDD SGR (pFK_i389_F-Luc_EI_NS3-3_JFH1_deltaGDD) has a deletion of the GDD motif within the viral polymerase. The p100 NS3-3’ UTR SGR was generated by inserting a fragment from p100NGS_coding into pFK_i389_F-Luc_EI_NS3-3_JFH1 using the restriction enzymes MreI and BbvCI.

All sequences of newly generated plasmids were confirmed by Sanger sequencing (Microsynth AG). More detailed information about the cloning strategy is available upon request.

### Cell Culture

Immortalized human hepatoma cell lines Huh-7 [80], Huh-7.5 [23, 81], Huh-7.5 miR-122 -/- [17], Hep-G2 hCD81, HuH6 hCLDN1 [13] and Lunet N#3 and Lunet N#3 hCD81 [18] were cultured in Dulbecco’s modified Eagle’s medium (DMEM) (Gibco, Thermo Fischer Scientific) supplemented with 1 % MEM non-essential amino acids (Gibco, Thermo Fischer Scientific), and 100 U/ml penicillin, 100 µg/ml streptomycin (Gibco, Thermo Fischer Scientific) and 2 mM L-glutamine (Gibco, Thermo Fischer Scientific) designated as DMEM complete. For routine culture DMEM complete was further complemented with 10 % fetal calf serum (FCS) (Capricorn Scientific). Culture medium of Hep-G2 hCD81 and HuH6 hCLDN1 cells was additional supplemented with 5 µg/ml blasticidine (Thermo Fischer Scientific). Additionally, Hep-G2 cells were cultured on collagen-coated plates (phosphate-buffered saline (PBS) containing 0.1 % acetic acid and 0.01 % collagen (Serva Electrophoresis GmbH)). Culturing was performed at 37 °C in 5 % CO_2_. All cell lines were regularly tested by standardized PCR service (Eurofins) for mycoplasma contamination.

The Primary Human Hepatocyte Core Facility (Hannover Medical School) isolated and seeded PHHs from liver explant tissue as previously described [82]. PHHs were maintained in HBM basal medium complemented with HCM SingleQuots (Lonza) and 100 U/ml penicillin, 100 µg/ml streptomycin (Gibco, Thermo Fischer Scientific). The experimental usage of PHHs is approved by the ethics commission of Hannover Medical School (vote no. 252-2008).

### *In vitro* transcription

HCV full-length genome or sub genome containing plasmid DNA (10 µg) was linearized according to the restriction enzyme recognition sequence located at the 3’ end of the viral genome. Linearized plasmid was purified using Qiaprep Spin Miniprep Kit (Qiagen). For T7-based *in vitro* transcription, 100 µl containing 2 µg plasmid DNA in 60 µl H_2_O, 20 µl 5x RRL buffer (1 M HEPES, pH 7.5, 1 M MgCl_2_, 1 M Spermidine, 1 M DTT in H_2_O), 12.5 µl rNTP solution (25 mM each rNTP), 100 U RNasin ribonuclease inhibitor (Promega) and 60 U T7 polymerase (Promega) were incubated at 37 °C for 2 h. *In vitro* transcription was boosted by freshly adding 30 U T7 polymerase for further 2 h of incubation time. *In vitro* transcription reaction was terminated by addition of 7.5 U RQ1 Dnase (Promega) to the reaction mix and incubation for 30 min at 37 °C. *In vitro* transcribed RNA was purified using the NucleoSpin RNA Clean-up kit (Macherey-Nagel). RNA concentration and purity was evaluated using the NanoDrop One (Thermo Scientific). Quality of the *in vitro* transcripts was checked by agarose gel electrophoresis.

### HCVcc production

For HCVcc production, 6 × 10^6^ Huh-7.5 cells in 400 µl Cytomix (120 mM KCl, 0.15 CaCl_2_, 10 mM KHP0_4_, 25 mM HEPES, 2 mM EGTA, 5 mM MgCl_2_, 2 mM ATP, 5 mM glutathione) were electroporated (270 V, 975 µF, cuvette gap width 0.4 cm) with 10 µg of *in vitro* transcribed HCV full-length RNA using the Gene Pulser Xcell system (BioRad). Cells of two electroporations were combined in 20 ml of DMEM complete and seeded into one 15 cm dish. On the next day, medium was exchanged to DMEM complete containing 2 % FCS, before cell culture supernatant was collected, filtered (0.45 µm) at 48, 72 and 96 h.p.e. Pooled, filtered cell culture supernatant containing infectious HCV was concentrated 50x using 100 kDa Amicon Ultra-15 centrifugal filter units (Merck Millipore) as recommended by the manufacturer. To prepare comparable virus stocks of the infectious clones and population, a 15 cm dish of naïve Huh-7.5 cells was infected using the concentrated virus stock and maintained for 3 days. After a second round of infection by passaging the infectious cell culture supernatant from the 15 cm dish of infected Huh-7.5 cells on four 15 cm dishes of fresh Huh-7.5 cells, medium was exchanged to DMEM complete supplemented with 2 % FCS at 24 hours post infection. Starting from 48 hours post infection, cell culture supernatant was again collected, filtered and pooled every 24 hours. The resulting virus stock was concentrated 25x as described earlier. The resulting highly infectious master HCV stocks were characterized by calculating the tissue culture infectious dose 50 per ml (TCID_50_/ml) upon infection of Huh-7.5 cells and measuring the containing genome equivalents (GE) per ml by an HCV-specific RT-qPCR.

### HCV infection assays

Huh-7 and its derivatives were seeded at a density of 5 × 10^4^ per well of a 24-well plate at a minimum of 16 hours prior to inoculation. For HuH6 hCLDN1 and Hep-G2 hCD81 cell lines, a density of 1 × 10^5^ and 1.4 × 10^5^ per well was chosen, respectively. For PHHs an approximal density of 3.75 × 10^5^ viable cells per well was aimed but varies depending on the cell quality after the isolation process. PHH were either cultured without or with 10 µM ruxolitinib pretreatment prior to infection. To analyze replication fitness of different HCV variants, cells were inoculated with normalized amounts of GE per cell as indicated in the corresponding figure legend. After 4 hours of incubation at 37 °C and 5 % CO_2_, the inoculum was aspirated, and residual unattached virus particles removed by washing of the cells for 4 times with PBS before fresh cell culture medium, including ruxolitinib where indicated, was applied. Replication fitness was analyzed by collection of supernatant and cell lysates at 4, 24, 48 and or 72 h.p.i. Collected supernatant was used to measure virus particle release over time by determination of the TCID_50_/ml titer. Total RNA was extracted from cell lysates and used to measure intracellular viral RNA levels by RT-qPCR.

### HCV replication assays

For replication assays, 5 µg of *in vitro* transcribed HCV SGR RNA was mixed with 6 × 10^6^ Huh-7 cells in cytomix solution and electroporated as described in detail in the HCVcc production protocol. Electroporated cells were transferred into DMEM complete supplemented with 10 % FCS, reaching a density of 3 × 10^5^ cells/ml. From this 3 × 10^4^ cells per well were seeded in quadruplicates in one 96-well plate per analyzed time point. At 4, 24, 48 and 72 hours post electroporation, cell culture supernatant was removed, and cells lysed directly on the plate by addition of 35 µl/well luciferase lysis buffer (1 % Triton X-100, 25 mM diglycine, 15 mM MgSO_4_, 4 mM EGTA, 1 mM DTT) and storage at -20 °C prior to luciferase measurement.

### HCV entry and assembly assays

To study HCV entry and assembly phenotypes of different HCV variants, 0.75 µg of *in vitro* transcribed HCV full length RNA was electroporated into 4 × 10^6^ Lunet N#3 cells similar to the HCVcc production protocol. Subsequently after the electroporation, Lunet N#3 cells were seeded in 6-well format in a density of 4 × 10^5^ cells per well. The kinetics of viral particles release upon transfection of HCV RNA was analyzed by collecting the cell culture supernatant of electroporated cells at 4, 24, 48 and 72 hours post electroporation. Virus titer of collected supernatants was measured using an end-point dilution assay to determine the TCID_50_/ml.

### HCV receptor blocking assay

Huh-7.5 cell were seeded in a density of 2 × 10^4^ cell per well of a poly-L-lysine-coated 96-well plate. Two days after seeding, cells were incubated with a serial dilution from 10 to 0.0006 µg/ml of either anti-CD81 (JS-81; 555675, BD Pharmingen), mouse IgG1 κ isotype control (107.3, 554721, BD Pharmingen) or anti-SR-BI (C16-71) [83]. After 1 hour of incubation at 37 °C, cells were infected with an amount of virus that results in approx. 200 FFU/well. After further incubation for 3 hours, one PBS wash was performed, and fresh media added. At 48 hours after infection, cells were fixed with ice-cold methanol and stained for NS5A expression by immunofluorescence staining.

### HCV spreading assay

To study cell-to-cell spread, 2 × 10^6^ Lunet N#3 cells were electroporated with 1.25 µg of *in vitro* transcribed HCV full-length RNA as stated earlier in the material and method section. Electroporated cells were transferred into 9.6 ml of DMEM complete containing 10 % FCS to reach a final concentration of 2 × 10^5^ cells per ml. Electroporated cells were further diluted in a cell suspension containing a similar cell density of naïve Lunet N#3 (1:50 dilution ratio) or Lunet N#3 hCD81 (1:100 dilution ratio). Undiluted and diluted cells were seeded in a density of 2 × 10^4^ cells/well into a 96-well plate and incubated for 48 hours. Cells were then fixed by incubation in 3 % paraformaldehyde (PFA) for 30 min before NS5A expression was detected using immunofluorescence staining.

### Drug sensitivity assay

Huh-7 cells were electroporated with 5 µg of *in vitro* transcribed SGR RNA as done in the HCV replication assay. Electroporated cells were seeded in half of a 96-well plate per drug with a density of 3 × 10^4^ cells per well. At 5 h.p.e., standard cell culture medium was exchanged to DMEM complete containing 10 % FCS, a constant end concentration of 0.01 % DMSO and serial dilutions of telaprevir (S1538, Selleckchem), daclatasvir (S1482, Selleckchem) and cyclosporin A (C988900, Toronto Research Chemicals). In the case of the Intron A solution (Merck Sharp & Dome GmbH), serial dilutions in DMEM complete containing 10 % FCS did not require the addition of DMSO as a solvent control. After 48 hours of incubation, cell culture supernatant was removed and cells lysed on the plate by addition of 35 µl/well of luciferase lysis buffer (1 % Triton X-100, 25 mM diglycine, 15 mM MgSO_4_, 4 mM EGTA, 1 mM DTT). Cell lysates were stored at -20 °C until luciferase measurement was performed.

### Virus titer determination

Virus titer of cell culture supernatant was determined by a limiting dilution assay followed by calculating the TCID_50_/ml after immunohistochemical staining of cells inoculated with the supernatant. For this, highly permissive Huh-7.5 cell were seeded at a density of 1 × 10^4^ cells in 200 µl per well of a 96-well plate. On the day after seeding, cells were inoculated in replicates of 6 wells per condition with serial dilutions of the cell culture supernatant. The dilution ratio (1:3, 1:5 or 1:10) was chosen depending on the estimated infectivity of the culture fluid. After 72 hours of incubation at 37 °C and 5 % CO_2_, cell culture fluid was removed, and cells fixed by addition of ice-cold methanol for 20 min. Cells were washed two times with PBS containing 0.1 % Triton X-100 (PBS-T). Blocking of endogenous peroxidase activity was performed by incubation with 3 % H_2_O_2_ in PBS-T for 5 min at RT. NS5A expression in infected cell was detected by incubation with mouse anti-NS5A monoclonal antibody 9E10 [15] (Cell essentials) in a concentration of 0.44 µg/ml in PBS-T for 1 hour at RT. Bound NS5A antibody was detected after two washes with PBS-T by incubation with horseradish peroxidase (HRP)-conjugated anti-mouse antibody (A4416, Sigma-Aldrich) diluted 1:300 in PBS-T. After 1 hours of incubation, cells were washed twice with PBS-T before filtered substrate solution (28.78 mM sodium acetate, 11.52 mM acetic acid, 4.42 mM 3-amino-9-ethylcarbazole in N, N-dimethylformamide, 0.3 % hydrogen peroxide) was added as a substrate of the HRP-conjugated secondary antibody. HRP activity was terminated after a minimum of 10 min incubation at room temperature by removing the substrate and addition of distilled H_2_0. TCID_50_/ml was determined by counting NS5A positive wells using the bright field microscope followed by the calculation using the TCID_50_ calculator [84].

### Immunofluorescence staining

After fixation for immunofluorescence staining, residual PFA was washed away using PBS. Cells were permeabilized for 5 min at RT using 0.2 % Triton X-100 in PBS. Residual Triton X-100 was washed away using PBS before blocking in 5 % horse serum in PBS for 1 hour. Then cells were incubated for 1 hour at room temperature (RT) with an anti-NS5A monoclonal antibody 9E10 [15] (Cell essentials) in a concentration of 0.44 µg/ml in PBS containing 5 % goat or horse serum. Bound NS5A antibody was detected after two washes with PBS by incubation with anti-mouse Alexa Fluor 488 plus (A32723, Thermo Scientific) or Alexa Fluor Plus 555 (A32773, Thermo Scientific) in a concentration of 2 µg/ml in 5 % goat or horse serum in PBS. After 1 hour of incubation, cells were washed twice with PBS before cellular DNA was stained by a short incubation with 4′, 6-diamidino-2-phenylindole (0.5 µg/ml). Lastly, three washes with water were performed prior to microscopic analyses. Microscopic images were taken using Immunospot analyzer (CTL Europe GmbH) or the Keyence BZX800 microscope (4x objective). The number of FFU were counted using the analyzing software of the Immunospot analyzer or FIJI in combination with CellProfiler [85, 86]. For visualization of full 96-well fluorescence images a maximum filter (r=10 pixels) was applied as previously described [87].

### RT-qPCR

Total RNA from liver cells was extracted using the NucleoSpin RNA kit (Macherey-Nagel). Viral RNA from cell culture supernatant was isolated using the Quick-RNA Viral kit (ZymoResearch). Both preparations were performed according to the recommendations of the manufacturer. The number of HCV genome equivalents present in the samples was measured by an HCV probe-based approach. For this 2 µl of RNA sample were mixed with the PCR ingredients of the one-step LightCycler 480 RNA Master Hydrolysis Probes kit (Roche Holding AG), the JFH1-specific probe (5’-6FAM-AAA GGA CCC AGT CTT CCC GGC AA-TMR-3’) as well as the primers S-146 (5’-TCT GCG GAA CCG GTG AGT A-3’) and A-221 (5’-GGG CAT AGA GTG GGT TTA TCC A-3’) according to manufacturer’s instructions. The PCR reaction protocol was set as follows: 63 °C for 3 min, 95 °C for 30 sec and 45 cycles with each 95 °C for 15 sec and 60 °C for 30 sec, before a final cooling 40 °C for 30 sec was performed. The HCV GE were determined using serially diluted *in vitro* transcribed HCV full-length RNA with a known copy number as an internal standard, followed by normalization to the total RNA of each sample. Technical duplicates were performed for each sample.

### Luciferase measurement

After thawing, 20 µl/well of cell lysates were transferred to a white luminometer 96-well plate containing 72 µl assay buffer (25 mM diglycine pH 7.8, 15 mM KPO_4_, 15 mM MgSO_4_, 4 mM EGTA pH 8, 1 mM DTT, 2 mM ATP pH 7.8) before 40 µl of luciferin solution (0.2 mM D-luciferin, 25 mM diglycine) were automatically added and bioluminescence measurement performed by the Berthold LB 960 microplate reader (Berthold Technologies).

### RNA sequencing and transcriptome analyses

For transcriptional profiling of HCV-infected cells, 4 × 10^5^ Huh-7 cells and 2 x 10^6^ PHHs were seeded per well on a 6-well plate. On the day of infection, cell culture medium was replaced by 1 ml of fresh medium before virus inoculum (MOI 1 TCID_50_/ml per cell) or an equal volume of conditioned cell culture medium from uninfected cells, which underwent the same preparation as the virus stock (e.g., passaging and concentration), was added. After 4 hours of incubation, the inoculum was topped up with 1 ml of fresh cell culture media. After further 24 hours of incubation, cell culture supernatant was completely removed, and cells washed once with PBS. Cells were lysed in 350 µl of RA1 buffer of the NucleoSpin RNA kit (Macherey-Nagel) supplemented with 1 % beta-mercaptoethanol followed by passing ten times through a narrow-bore syringe (Omican-F, 1 ml, 0.3 × 12 mm, 9161502, B Braun). Total RNA was extracted according to manufacturer’s instructions. Quality and quantity of extracted RNA was analyzed using the NanoDrop One (Thermo Scientific) and Fragment Analyzer (Agilent). Library preparation was performed according to NNSR priming principle [88] as described previously [8]. The resulting libraries were sequenced using the NextSeq 550 platform (Illumina) with single end 1x 86 bp setting. After trimming the two initial nucleotides from the reads, the transcriptome analyses were performed using CLC Genomics Workbench Version 21.0.4 (Qiagen). Raw FASTQ files were aligned against the human reference genome GRCh38 (updated on the 24.11.2021). Identification of differentially expressed genes was done by filtering on average expression for false discovery rate (FDR) correction and testing differentially expression due to infection while controlling for passage number of Huh-7 cells or the donor of the PHHs. Further Gene Ontology (GO) enrichment analysis settings included exclusion of data with a mean RPKM below 0.5 and a fold change below 1.5 and a threshold of FDR p-value > 0.05. Further processing of Gene Set Enrichment and KEGG pathway analyses were performed in R [89] using the gseKEGG function from the R package ‘clusterProfiler’ v4.7.1.001 [90]. Volcano plots were generated using the R package ‘EnhancedVolcano’ v1.16.0 [91]. PCA, volcano, GO and KEGG plots were all generated or improved with the R package ‘ggplot2’ v3.4.1 [92].

### Statistical analyses

Experiments were conducted in a minimum of three independent biological replicates. Graphical visualization of experimental data and statistical analyses were performed with GraphPad Prism 9 (GraphPad Software). Prior to statistical analyses, data were transformed into logarithmic format (not done for Fig. 6J). Statistical significance was then calculated using either two-tailed, unpaired t-test or two-way ANOVA with either Sidak’s or Dunnett’s multiple comparisons correction as indicated in the respective figure legend. Significance levels were visualized as follows: p < 0.05 = *, p < 0.01 = **, p < 0.001 = *** and p < 0.0001 = ****.

### Data availability

The raw RNA-seq data and subsequent downstream analyses (gene expression browser, GO enrichment analysis) were submitted to the NCBI GEO database and can be accessed under the accession number GSE246981.

## Acknowledgements

Huh-7.5 cells were kindly donated by Charles M. Rice (Rockefeller University, New York City, USA). Matthew Evans (Icahn School of Medicine at Mount Sinai, New York, USA) kindly provided the Huh-7.5 miR-122 -/- and the corresponding parental cell line. Furthermore, p100pop was a kind gift of Esteban Domingo (CBMSO, Madrid, Spain) and we appreciate the fruitful project discussion with him and his team. The JFH1 subgenomic replicon plasmids were kindly provided by Volker Lohmann (Heidelberg University, Heidelberg, Germany).

## Author contributions

N.F.: conceptualization, data curation, formal analysis, investigation, visualization, writing – original draft preparation, writing – review and editing

R.J.P.B.: data curation, formal analysis, resources, software, visualization, supervision writing – original draft preparation

B.M.R.: investigation, data curation, formal analysis, software, writing – review and editing

M.H.: formal analysis, software, data visualization, writing – review and editing

Y.B.: formal analysis, software, visualization, writing – review and editing

D.T.: formal analysis, visualization, writing-review and editing

C.M.: formal analysis, software, data curations, writing – review and editing

F.W.R.V.: resources, writing – review and editing

E.S.: resources, supervision, writing – review and editing

T.P.: conceptualization, funding acquisition, methodology, resources, supervision, writing – review and editing

J.S.: conceptualization, data curation, formal analysis, funding acquisition, investigation, methodology, project administration, resources, supervision, writing – original draft preparation, writing – review and editing

